# Towards population genetic assessments and species abundance from environmental DNA: A case study with zebrafish in controlled aquaria

**DOI:** 10.1101/2025.02.15.638407

**Authors:** Abinaya Meenakshisundaram, Simon Jarman, Haylea Power, Jason W. Kennington, Luke Thomas

## Abstract

Developing robust methods for amplifying and analysing highly-polymorphic nuclear genetic markers from environmental samples could assist in the reliable and scalable long-term monitoring of elusive, threatened or invasive species that are otherwise challenging to observe. In this study, we used zebrafish in controlled aquaria to apply forensic science approaches and demonstrate that microhaplotypes, which are short segments of nuclear DNA (100–300bp) containing two or more single nucleotide polymorphisms (SNPs), can be amplified from trace DNA in water samples to accurately estimate population genetic diversity and species abundance. We successfully amplified a panel of 17 microhaplotypes that comprised 69 SNPs which could reliably estimate population-level allele frequencies and genetic diversity estimates from water DNA. The panel of microhaplotypes amplified from water samples from replicate tanks strongly matched allele frequency estimates from corresponding tissue samples, and could also be used for estimating number of contributors from multi-individual samples. Our research demonstrates the effectiveness and potential of amplifying microhaplotype panels from eDNA as a non-invasive and scalable tool for population genetic studies of aquatic species.

## 1. Introduction

Population genetic monitoring of wildlife species is currently used to study changes in genetic diversity, population structure, connectivity, abundance, adaptation and fitness of populations. This may help to inform on appropriate management strategies for some species in changing environments (Crawford & Oleksiak, 2016; Reed & Frankham, 2003; Von Der Heyden et al., 2014; Yi et al., 2023). Measurement of abundance from genetic monitoring is currently limited to studies where individual samples can be collected directly (tissue biopsy, fin clippings, etc) (Pirog et al., 2019; Vignaud et al., 2014) or via non-invasive sampling techniques (scats, shed skin, parasites, etc) (Meekan et al., 2017; Parsons et al., 1999; Pierszalowski et al., 2013).

DNA traces left by organisms in the environment, referred to collectively as environmental DNA (eDNA), can be used to gather important demographic and ecological data for areas and species of conservation importance (Taberlet et al., 2018). In the last decade, eDNA from marine environments has been increasingly applied to include ecosystem-wide estimates of biodiversity to single species detection (Adrian-Kalchhauser & Burkhardt-Holm, 2016; Berry et al., 2019; Dugal et al., 2022b; Dugal et al., 2023; Nevers et al., 2018). However, its application to study population genetics of species is much more limited, but shows great promise for assessing genetic diversity, population structure, and demographic patterns without direct animal interaction (Adams et al., 2019; Foote et al., 2012; Suarez-Bregua et al., 2022). This approach can be particularly advantageous for genetic sampling of cryptic, elusive or highly migratory marine species that inhabit expansive, remote or hard-to-access habitats, or that tend to traverse large distances and spend major parts of their life cycle in deep open waters. By using genetic markers found in water samples, eDNA has the potential to provide population-level insights of an aquatic species, while potentially reducing costs, effort and risks to researchers associated with traditional methods like tagging or visual surveys which require closer interaction, as well as ethical challenges that might arise in capturing or invasively sampling a protected species (Adams et al., 2019; Foote et al., 2012).

Previous research using eDNA for population genetic studies have focused mainly on short, highly polymorphic regions of the mitochondrial DNA due to the high copy number and thus higher probability of detection in water samples (Adams et al., 2024; Dejean et al., 2011; Duarte et al., 2023; Dugal et al., 2022a). However, nuclear DNA markers offer greater resolution and a wider range of evolutionary insights such as demographic trends, gene flow, effective population size, species abundance, and the ability to distinguish individuals from a mixed DNA sample (Adams et al., 2019; Andres et al., 2023; Suarez-Bregua et al., 2022). Advancements in human forensic science have addressed the challenge of identifying individuals from mixed DNA samples found in crime scenes through the use of short, multiallelic segments of nuclear DNA called “microhaplotypes (MH)” (Kidd et al., 2013; Oldoni et al., 2019). These multiallelic loci of approximately 100–300 bp containing two or more single nucleotide polymorphisms (SNPs) (Kidd et al., 2014) can be sequenced in a single run and provide important forensic data regarding individual detection, biogeographic ancestry and lineage / familial relationships (Chen et al., 2018a; Oldoni et al., 2017, 2019, 2020). MH marker panels have been used successfully to monitor wild terrestrial and marine animal populations through direct invasive genetic sampling of individuals for studying population structure (Baetscher et al., 2022; Hopken et al., 2023), familial relationships (Baetscher et al., 2018), gene flow and potential local adaptations (Morin et al., 2021). Although MH markers show dramatically increased power and efficiency in direct genetic samples compared to di-allelic SNP markers and eliminates the problem of stutter or allelic dropout that is associated with microsatellite markers (Andres et al., 2023), its applicability in eDNA samples has not been explored.

In this study, we used zebrafish (*Danio rerio*) as a model system in a controlled experimental aquaria environment to study the efficiency and limitations of estimating population-level genetic diversity statistics and the number of unique individuals in complex water samples using a novel panel of species-specific MH markers developed by adopting forensic science-based techniques. This framework would provide a basis for developing eDNA population genetic monitoring techniques for other marine and freshwater species.

## 2. Methodology

### 2.1 Sample collection & processing

411 zebrafish individuals (AB strain (wild-type, ZIRC catalogue ID ZL1)) aged 3-5 months were maintained in 3.5 L standard aerated tanks by Animal Care Services at the Zebrafish Experimental Research Centre, Shenton Park, Western Australia. Animal care, handling, collection and use of samples were done with approval from Animal Ethics committee of the University of Western Australia (application number: RA/3/100/1732). The zebrafish were kept in four tanks of 2, 4, 10, 20 individuals each and 15 tanks containing 25 individuals each, labelled as 25(1), 25(2), 25(3), … 25(15), with equal numbers of males and females when possible. Prior to collection of water samples for the purpose of extracting eDNA, the zebrafish were transferred from their aerated tanks into stagnant water tanks where they were held for 3 hours before transfer back to housing tanks. The zebrafish groups with 2, 4, 10 and 20 individuals were placed in 3 L and groups with 25 individuals were held in 8 L tanks of stagnant UV treated and filtered water. Prior to placement of zebrafish in the tanks, 1 L of the UV treated filtered water was taken as negative control to check for DNA contamination. After the 3-hour holding period, the water in the tanks were homogenized by manual stirring and 1 L of water samples were collected from each tank in Nalgene bottles sterilized using 10% bleach solution, 70% ethanol, followed by rinsing in deionised (DI) water, resulting in eDNA samples comprising of 2, 4, 10, 20 and 25 genetic contributors. 1 L of the remaining water from each tank were pooled in combinations shown in Table 1 to make up eDNA samples containing higher number of contributors of 50, 100, 200, 300 and 411 individuals. The pooled water samples were homogenized by manual stirring and 1 L of water from each pool were collected using sterile Nalgene bottles. This process of eDNA collection was repeated three times on separate days. The water samples were immediately transported (within 2-3 hours) to a molecular biology laboratory (Indian Ocean Marine Research Centre (IOMRC), The University of Western Australia) on ice. The samples were filtered in a sterile laminar airflow hood using sterile single use 250 ml EZ-fit funnel filtration units with 0.22 μm filter membranes and an EZ-fit Manifold base from Merck Millipore (Merck Group, www.sigmaaldrich.com) (Kumar et al., 2020; Majaneva et al., 2018). The work space and manifold base were sterilized using 10% bleach solution followed by 70% ethanol solution between each sample. Filter papers containing the eDNA samples were stored at -80 °C in cryogenic tubes. DNA was extracted from half of the filter papers using the DNeasy Blood & Tissue kit (QIAGEN www.qiagen.com) following manufacturer’s protocol with the modification of adding 360 μL ATL buffer and 40 μL proteinase K for overnight incubation at 56 °C (Dugal et al., 2022a). The supernatant obtained was used for automated DNA extraction using a QIAcube DNA extraction robot (QIAGEN) to prevent cross contamination between samples due to human errors. The DNA samples were stored at -80 °C until further use.

**Table 1:** The number of contributors in pooled water samples from zebrafish tanks. Samples were taken from four tanks containing 2, 4, 10, or 20 zebrafish, as well as from 15 separate tanks with 25 zebrafish each (labelled 25(1), 25(2), 25(3), …, 25(15)) and pooled using combinations outlined in the column constituents.

| Sample name | Constituents | Volume of pool (L) | Volume filtered (L) | No. of contributors |
| --- | --- | --- | --- | --- |
| 50(1) | 25(1), 25(2) | 2 | 1 | 50 |
| 50(2) | 25(2), 25(3) | 2 | 1 | 50 |
| 100(1) | 25(1), 25(2), 25(3), 25(4) | 4 | 1 | 100 |
| 100(2) | 25(4), 25(5), 25(6), 25(7) | 4 | 1 | 100 |
| 100(3) | 25(8), 25(9), 25(10), 25(11) | 4 | 1 | 100 |
| 100(4) | 25(12), 25(13), 25(14), 25(15) | 4 | 1 | 100 |
| 200(1) | 25(1), 25(2), 25(3), 25(4), 25(5), 25(6), 25(7), 25(8) | 8 | 1 | 200 |
| 200(2) | 25(4), 25(5), 25(6), 25(7), 25(8), 25(9), 25(10), 25(11) | 8 | 1 | 200 |
| 200(3) | 25(8), 25(9), 25(10), 25(11), 25(12), 25(13), 25(14), 25(15) | 8 | 1 | 200 |
| 300(1) | 25(1), 25(2), 25(3), 25(4), 25(5), 25(6), 25(7), 25(8), 25(9), 25(10), 25(11), 25(12) | 12 | 1 | 300 |
| 300(2) | 25(4), 25(5), 25(6), 25(7), 25(8), 25(9), 25(10), 25(11), 25(12), 25(13), 25(14), 25(15) | 12 | 1 | 300 |
| 411 | 2, 4, 10, 20, 25(1), 25(2), 25(3), 25(4), 25(5), 25(6), 25(7), 25(8), 25(9), 25(10), 25(11), 25(12), 25(13), 25(14), 25(15) | 19 | 1 | 411 |

After collection of the last replicate of eDNA samples, all zebrafish were euthanized using an ice slurry and immediately transported to the molecular biology lab for further processing. Tissue samples in the form of tail and/or fin clippings sufficient for DNA extraction were collected from the euthanized individuals while maintaining sterility and preventing cross contamination between samples. The tissue samples were extracted for DNA using the DNeasy Blood & Tissue kit following manufacturer’s protocol and stored at -80 °C until use. All handling of tissue, eDNA samples and DNA extractions were performed in a laboratory separate from where PCR reactions and amplicons were handled to limit contamination (Goldberg et al., 2016).

### 2.2 Characterizing microhaplotypes

To develop a panel of MH markers, DNA samples from 96 zebrafish individuals (AB strain (wild-type, ZIRC catalogue ID ZL1)) sourced from Western Australian Zebrafish Facility at the University of Western Australia were sent to Diversity Arrays Technology Pty Ltd (www.diversityarrays.com) for DArTseq to detect allelic variations from several hundred loci across the genome (Kilian et al., 2012). Mixture detection and deconvolution study on human DNA MH markers revealed a high probability of mixture detection (95%) using as few as five MH markers with average effective alleles equal to or slightly above 3 and individual SNPs showing high heterozygosity (Kidd et al., 2014; Kidd & Speed, 2015). Using these criteria, SNP data across 21995 loci acquired from DArTseq were filtered to find target sequences containing two or more SNPs with individual SNP heterozygosity > 18%. Heterozygosities were estimated using dartR package on R (Gruber et al., 2018) and a total of 78 sequences of length 70 bp each passed our filtration. The target sequences found were aligned to the reference genome GRCz11 (Genome Reference Consortium) using ‘blastn’ function of the National Center for Biotechnology Information (NCBI) BLAST version 2.2 (Camacho et al., 2009). Flanking regions of 100 bp on either side of the aligned sequences were extracted from the reference genome using BEDtools v2.31.0 (Quinlan, 2014). These extracted sequences were aligned against the entire ‘RefSeq Genome Database’ of blastn (NCBI BLAST) to check for species specificity as well as against the zebrafish reference genome to check for sequence specificity (Johnson et al., 2008). Primers were designed for the selected MH sequences using primer-BLAST (NCBI) (Ye et al., 2012) with melting temperature (*T_m_*) 55 ± 5 °C, primer length between 18 – 30 bp, GC% from 40 - 60%, resulting amplicon lengths from 150 to 270 bp and for sequence specificity within the zebrafish genome. Multiplex compatibility of the primer pairs was checked using PrimerPlex 2.76 (PREMIER Biosoft, http://www.premierbiosoft.com) (Kechin et al., 2020). As a result, a set of 50 primer pairs potentially amplifying a panel of newly characterized MH markers were developed and ordered from Alpha DNA (http://www.alphadna.com) after the inclusion of an Illumina overhang adapter sequence.

### 2.3 Optimizing multiplex assay

All 50 newly designed primer pairs were individually tested using zebrafish tissue DNA for presence of target and non-target amplifications. The individual test PCR reactions of 10 μL were set up with 5 μL of HotStarTaq Master Mix (QIAGEN), 0.2 μM each of the corresponding forward and reverse primers and 2 ng tissue DNA. A two-step Touch-Down PCR (TD-PCR) was performed by introducing an initial set of cycles at varying primer annealing temperatures prior to the basic PCR cycles to ensure binding of all primer pairs to their respective target sequences (Moezi et al., 2019). The thermocycler conditions were: 95 °C for 15 minutes, 15 cycles of denaturation at 95 °C for 30 seconds, annealing at 60 °C with negative 1 °C decrements every cycle for 30 seconds, elongation at 72 °C for 1 minute, followed by 30 basic PCR cycles of 95 °C for 30 seconds, annealing at 56 °C for 30 seconds, elongation at 72 °C for 1 minute, and a single elongation step at 72 °C for 10 minutes. The PCR products were checked on a LabChip GX Touch HT fragment analyser using a DNA high sensitivity assay kit (Perkin Elmer, content.perkinelmer.com). 40 primer pairs out of the 50 showed amplification of MH amplicons at expected fragment sizes without non-target amplifications.

Optimal multiplex PCR assay to amplify all 40 target MH markers through a single reaction was developed with each 25 μL reaction volume containing 12.5 μL of 2x QIAGEN Multiplex PCR Master Mix (QIAGEN), 0.02 μM of each of the 40 primer pairs and 2 ng of tissue DNA or 5 μL of eDNA which contained DNA in concentrations ranging from 0.5 – 3 ng/μL. The thermocycler conditions for Multiplex TD-PCR were: 95 °C for 15 minutes, 15 cycles of denaturation at 95 °C for 30 seconds, annealing at 60 °C with negative 1 °C decrements every cycle for 90 seconds, elongation at 72 °C for 1 minute, followed by 40 basic PCR cycles of 95 °C for 30 seconds, annealing at 56 °C for 90 seconds, elongation at 72 °C for 1 minute, and a single elongation step at 72 °C for 10 minutes. The Multiplex PCR products were checked using the LabChip GX Touch HT nucleic acid analyser for presence of target fragment sizes (Supplementary Figure S1).

### 2.4 Library preparation for sequencing

Equal concentrations of extracted tissue DNA were made into artificially created pooled DNA mixtures with known number of contributors. The mixtures contained 2, 3, 4, 5, 6, 8, 10 or 20 individuals and were used for evaluating the efficiency of the genetic markers and optimizing the bioinformatic protocol to estimate the number of individual contributors to the DNA sample. To make the mixed samples, 1 ng of DNA from each individual sample was added to the mixture and 2 ng of this homogenized mixture was used to amplify MH markers for further analysis.

All DNA samples were amplified using the optimized multiplex PCR protocol to amplify the panel of 40 MH markers. The overall method involved a two-step PCR amplification protocol, with indexing of unique samples performed in the second step. The multiplex PCR products for each sample were indexed with different combinations of Illumina Nextera XT indices (Illumina www.illumina.com) for sample identification purposes. Index PCR reactions of 25 μL were set up using a QIAgility robotic workstation for prevention of cross-contamination or human error, with 12.5 μL of HotStarTaq Master Mix (QIAGEN), 2.5 μL each of the forward and reverse indices and 2 µL of multiplex PCR product if sourced from extracted tissue samples (including artificially pooled DNA mixtures) or 5 µL if sourced from extracted eDNA samples, due to the differences in starting DNA concentrations. Thermocycler conditions were: 95 °C for 15 minutes, 12 cycles of denaturation at 95 °C for 30 seconds, annealing at 55 °C for 30 seconds, elongation at 72 °C for 30 seconds, followed by a single elongation step at 72 °C for 10 minutes.

The PCR products were pooled at equimolar concentrations measured using a Qubit high sensitivity dsDNA kit (Thermo Fisher Scientific) to create library pools. For each PCR plate containing 96 samples, the samples were pooled into eight groups of 12 samples each for optimal removal of primer dimers and non-target fragments present in the product through size selection using 0.8X Ampure XP (Beckman Coulter www.beckmancoulter.com) magnetic bead clean-up. These pools were combined at equimolar concentrations to form the final library pool, which was checked using the LabChip GX Touch HT nucleic acid analyser for presence of target fragment sizes (Supplementary Figure S1). The final library was submitted to Genomics WA Laboratory in Perth, Australia (www.genomicswa.com.au) for sequencing using MiSeq Nano Kit V2 (300 cycle) (Illumina). Negative controls included during each stage (sample collection, DNA extraction and PCR) were checked for amplification using the LabChip nucleic acid analyzer after multiplex PCR amplification, and any showing amplification were sequenced following the same protocol as the samples.

### 2.5 Bioinformatic data processing

Sequence data received as demultiplexed fastq files were pre-processed using fastp-0.23.4 to perform quality control, remove low quality bases, trim adapters and poly-G tails (Chen et al., 2018b). The forward and reverse paired-end reads that passed initial QC filter were merged together using USEARCH v11, with maximum of five mismatches and minimum 90% alignment (Edgar, 2010). Consensus sequences of each of the expected 40 microhaplotype amplicon sequences was used as the reference to which the sequenced reads were mapped (Supplementary File S1). The reference amplicon sequences compiled in the form of a fasta file were indexed using samtools-1.12 (Li et al., 2009). BAM files of the merged reads mapped to the reference sequences were obtained using BWA-MEM (Li, 2013), filtered for mapping quality > 10, sorted and indexed using samtools-1.12 (Li et al., 2009). The number of sequences aligned to each MH amplicon reference sequence for each sample was calculated using samtools ‘idxstats’. Out of the panel of 40 MH markers, 17 (Supplementary Table S1) had a mean count of 50 or more sequences aligned to each reference amplicon sequence across both tissue and eDNA samples and were selected for further analyses.

SNP variants were called from the resulting BAM files using bcftools-1.3.1 (Danecek et al., 2021). Bcftools ‘mpileup’ and ‘call’ functions were used to exclude indels and call SNPs with an average depth across samples of > 10. The SNP sites were further filtered using vcftools 0.1.16 (Danecek et al., 2011) for Hardy-Weinberg Equilibrium (HWE) p-value ≥ 0.05. As a result, 69 SNP sites across 11 out of the 17 microhaplotypes were found common to both tissue and eDNA sample sequences. Probability of identity (*P_ID_*) as well as probability of identity between siblings (*PI_sibs_*) were calculated for all tissue-derived samples using function ‘pid_calc’ from R package PopGenUtils to evaluate the efficiency of this set of MH markers for individual identification (Taberlet & Luikart, 1999).

All subsequent analyses were performed using R version 4.3.1 (R Core Team, 2022). In all samples, alleles with less than 10 reads were removed to filter out sequencing or PCR errors (Andres et al., 2021). For eDNA samples, the read counts were scaled to 100 reads per sample to account for differences in starting DNA concentration and sequence read depths. Alleles with less than 0.01 scaled reads were removed. Scaled reads from the three replicate eDNA samples were pooled together due to low variation in read counts and allele frequencies between them. Allele frequencies were estimated from sequence read frequencies of alleles present in each sample as well as in the overall tissue or eDNA data set.

### 2.6 Contributor estimation using artificially made pooled DNA mixtures

Sequence data obtained from artificial pooled DNA mixtures derived from tissue, using up to 20 known individual contributors were used to validate the methodology for estimating individual and population level allele frequencies from mixed DNA samples, as well as to optimize bioinformatic methodology for estimating the number of individual contributors.

To assess accuracy of population allele frequency estimates, allele frequencies of the most common allele for each locus across all mixed samples were compared against the allele frequencies based on all individual tissue samples that contributed to the mixtures. The correlation between these frequencies was evaluated using Spearman’s correlation statistic through the ‘cor.test’ function of R package stats (R Core Team, 2022; Schober et al., 2018). The similarity between allele frequencies obtained from each mixed DNA sample and the pooled allele frequencies from the respective individual contributors were visualized using principal components analysis (PCA). All monomorphic loci and allele frequencies below 1% were removed prior to performing PCA using ‘prcomp’ function of R package stats.

To estimate the number of genetic contributors in a mixed sample, a “Likelihood-based mixture model” (Egeland et al., 2003; Sethi et al., 2019) was used which was developed in the field of forensics to provide likelihood estimates of the number of contributors (x) using the number (n) of observed alleles (A = {*a_1_,…,a_n_*}) and associated population allele frequencies (*p =* {*p_1_,…,p_n_*}) at each locus (*j*) as: 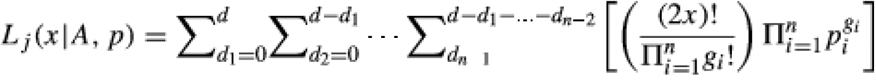 (Sethi et al., 2019) The maximum likelihood estimate of the number of contributors was given as the product of this likelihood across all loci. This model accounted for redundancy of alleles among individuals as well as masked alleles (*di*) that might be present in the mixture.

A custom R-script made by Andres et al. (2021) to obtain maximum-likelihood estimates of number of contributors using this model from eDNA samples using microsatellite markers was modified towards our purpose of obtaining contributor estimation using SNP data. Allele frequencies based on the entire 411 individual tissue data set from this study and those of the 38 individual tissue samples that contributed to the artificial pooled mixed DNA samples were taken as the population allele frequencies when estimating the number of individual contributors. For these analyses the data were filtered using increasingly strict thresholds of allele frequencies (0.001, 0.01, 0.1), and all alleles below this range were removed along with monomorphic loci prior to estimating the number of contributors (Andres et al., 2021). This was done to evaluate the sensitivity of this model to false alleles as well as the optimal filtering thresholds to be used to apply this estimation towards eDNA samples. The putative number of individuals (*y*) provided to the model was equal to the largest number of individuals present in a sample in the data set being analysed (*y* = 20). Additionally, to explore the efficiency of this model in estimating the number of contributors in mixed DNA samples versus individual tissue sample derived sequence data, we also estimated number of contributors using the combined allele frequencies of corresponding individual tissue samples that were the true contributors of each mixture.

### 2.7 Correlations between tissue and eDNA derived estimates of genetic diversity

The similarity between allele frequencies obtained from each eDNA sample and the combined allele frequencies from corresponding individual tissue samples that contributed to each sample were examined using a dendrogram based on Euclidean genetic distances generated using ‘dist’ and hierarchically clustered using the R package stats-4.3.1 (R Core Team, 2022) after removal of all monomorphic loci and alleles below 1% frequency.

Comparison of population allele frequencies of the most common allele for each locus from tissue versus eDNA samples was used to confirm the accuracy of estimates obtained from eDNA. The relationship between the population allele frequencies across all 69 loci calculated using all 411 individual tissue samples and all eDNA samples obtained in this study were evaluated using Spearman’s correlation statistic with the ‘cor.test’ function of R (R Core Team, 2022). Correlations between population allele frequencies based on all 411 tissue samples and those based on all three replicates of one single eDNA sample containing all 411 contributors, as well as all three replicates of eDNA samples containing 2, 4, 10, 20 and 25 individuals, with each of the 411 individual contributors being represented once were also evaluated.

Estimates of genetic diversity in the form of expected heterozygosity were calculated using the function ‘gl.report.heterozygosity’ with the R package dartR (Gruber et al., 2018) and adjusted for the corresponding sample size to obtain unbiased heterozygosity (uHe) values. Allelic richness (AR) were calculated using ‘allel.rich’ function of package PopGenReport (Adamack & Gruber, 2014). uHe and AR estimates to represent the entire population of individuals used in this study were based on SNP data from all 411 tissue samples, three replicates of a single eDNA sample containing all 411 individual contributors, and all three replicates of eDNA samples containing 2, 4, 10, 20 and 25 individuals, representing each of the 411 individual contributors once, without pooling of water samples. A paired Wilcoxon signed rank exact test implement using the ‘wilcox.test’ function of R (R Core Team, 2022) was used to test for differences in genetic diversity estimates obtained using the different methods.

Similarly, uHe and AR estimates were obtained from all three replicate eDNA samples from all 15 tanks containing 25 individuals each and compared it to the corresponding 25 tissue samples present in each tank. This was done to mimic eDNA samples obtained from field-based sampling and assess the ability to obtained accurate representations of genetic diversity of the contributors of each sample. Wilcoxon signed rank exact test was used as described earlier to test for the differences between genetic diversity estimates obtained from eDNA and tissue sampling methods.

### 2.8 Contributor estimation using eDNA

The R-script and optimized filtration parameters determined using the artificially made pooled DNA mixtures was used to estimate the number of contributors in each eDNA sample. Population allele frequencies from the entire tissue data set (411 individuals) were in the estimation of abundance in eDNA samples. All alleles with frequencies less than 1% were removed from each eDNA sample prior to analysis. The putative number of individuals (*y*) provided to the model was varied from 50, 411 or 1000 to observe its effect on contributor estimations.

## 3. Results

### 3.1 Sequence data

High read depth per sample was observed in zebrafish tissue samples (11875 (± 2631 SD)) and eDNA samples (12647 (± 2716 SD)) across all 11 microhaplotype markers (Supplementary Figure S2). The SNPs in tissue samples were found to have mean depth per site of 172 (± 68 SD) (Supplementary Figure S2) and average per-SNP quality of 56210.5 (± 44150 SD). Those called from eDNA sequences had a mean depth of 183 (± 69 SD) per site (Supplementary Figure S2) and average per-SNP quality of 12966.5 (± 10280.2 SD) showing high confidence in variant calls. The control sample consisting of UV treated filtered water from the zebrafish facility which was filtered and extracted similar to other eDNA samples did not show any sequenced reads after all loci with read counts below 10 was removed, except for one MH marker 52197511 which had a minimal number of reads (50 reads on average, which is 25% of that observed in other samples). All other negative controls for DNA extractions and PCR controls showed no amplification of MH markers. The combined probability of identity (*P_ID_*) for the 69 loci across all 411 individual tissue samples was estimated to be 1.15e-05 and *PI_sibs_* 0.0036, demonstrating the utility of this MH panel for individual identification.

### 3.2 Method validation using artificially created pooled DNA mixtures

Population allele frequencies obtained from all artificially pooled DNA mixtures were significantly correlated (ρ = 0.65 and *P* < 0.0001) to the combined allele frequencies of all corresponding individual samples (Supplementary Figure S3). Similarly, the PCA plot showed close grouping of samples based on allele frequencies of each pooled DNA mixture with the combined allele frequencies of the corresponding individual contributors (Supplementary Figure S3).

Contributor estimation using a maximum likelihood model on pooled DNA mixtures accurately estimated the number of contributors for mixtures of up to 10 individuals (Supplementary Figure S3). Using population allele frequencies from a larger dataset (411 samples) in the estimation reduced negative bias by 1-2 individuals compared to a smaller dataset (38 samples). Filtering alleles with frequencies below 0.001 or 0.01 yielded estimates closest to the true number of contributors, with a maximum bias of ±4 individuals for mixtures of up to 10 individuals, with accurate estimates obtained for mixtures with 4 and 5 contributors (Supplementary Figure S3). Filtering alleles with frequencies below 0.1 introduced higher negative bias, while retaining all alleles without frequency filtering led to high positive bias due to false positives or errors (Supplementary Figure S3). Retaining alleles with frequencies above 0.01 was chosen for eDNA analysis. Furthermore, all contributor estimates obtained from DNA mixtures and the combined allele frequencies of the corresponding contributing individual tissue samples were 100% similar.

### 3.3 Comparison between tissue and eDNA derived genetic data

The dendrogram constructed using Euclidean genetic distances based on allele frequencies showed high similarity between estimates based on tissue and eDNA samples, with each eDNA sample clustering close to the respective constituent individual tissue samples (Figure 1). Population allele frequencies obtained from all eDNA samples were highly correlated with associated tissue samples (ρ = 0.92, *P* < 0.0001, Figure 1). Similarly, strong associations were observed when population allele frequencies were calculated from the single eDNA samples containing all 411 contributors as well as when population allele frequencies were calculated from all the eDNA samples that represented each of the 411 individuals once in smaller batches of up to 25 individuals each (ρ = 0.86 and ρ = 0.93 respectively).

**Figure 1:**
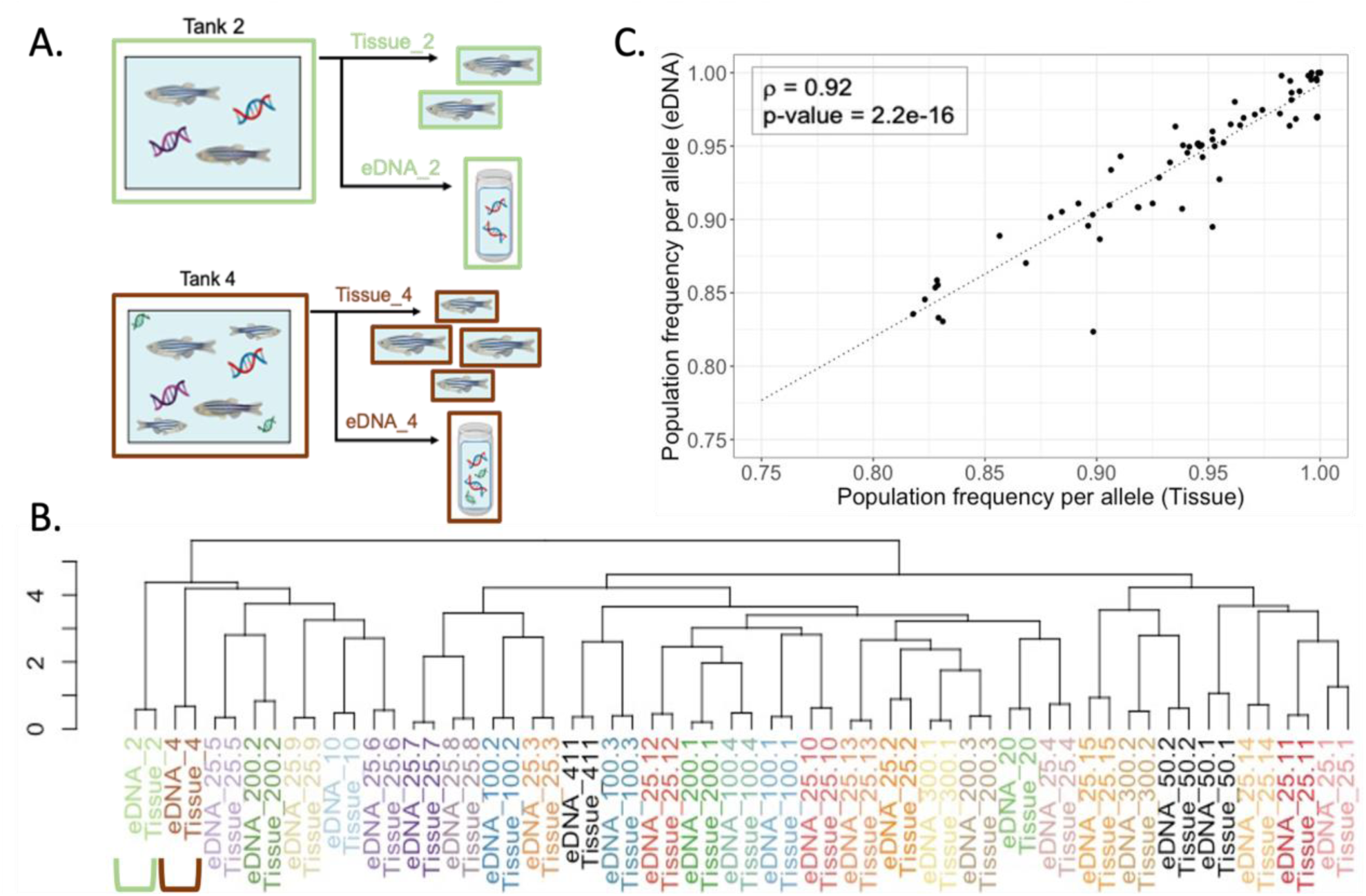
Comparative analysis of allele frequencies between environmental DNA (eDNA) and tissue samples from zebrafish tanks. (A) Schematic representation of the experimental workflow, illustrating sample collection from zebrafish tanks 2 and 4. Tissue samples were obtained from individual fish, while eDNA was collected from the tank water. Corresponding samples from each tank show strongly similar allele frequencies in the dendrogram below; (B) Dendrogram illustrating the hierarchical clustering of samples based on allele frequencies for eDNA samples and the combined allele frequencies from tissue samples of individual contributors in each corresponding mixture. Sample labels indicate the sample source, number of contributors (following the underscore) and the tank number (following the period); (C) Scatter plot showing relationship between population allele frequencies obtained across all eDNA samples and all tissue samples for the major allele at each SNP, with statistical significance of their correlation denoted by Spearman’s coefficient (ρ).

Estimates of expected heterozygosity obtained from each zebrafish tank containing 25 individuals each showed no significant differences between estimates obtained from eDNA samples and their respective tissue samples (p-value = 0.33, Figure 2). However, AR estimates obtained from eDNA samples in these tanks were significantly lower than those obtained from tissue samples (p-value = 0.00006, Figure 2). Whereas, overall population genetic diversity estimates (uHe and AR) from all 411 tissue samples showed no significant differences compared to eDNA samples from the same individuals (p = 1 and 0.19, respectively; Supplementary Figure S4). Estimates from a single eDNA sample with all 411 contributors were slightly lower, while those from three eDNA replicates, representing smaller groups but collectively all 411 individuals, closely matched tissue-based estimates (Supplementary Figure S4).

**Figure 2:**
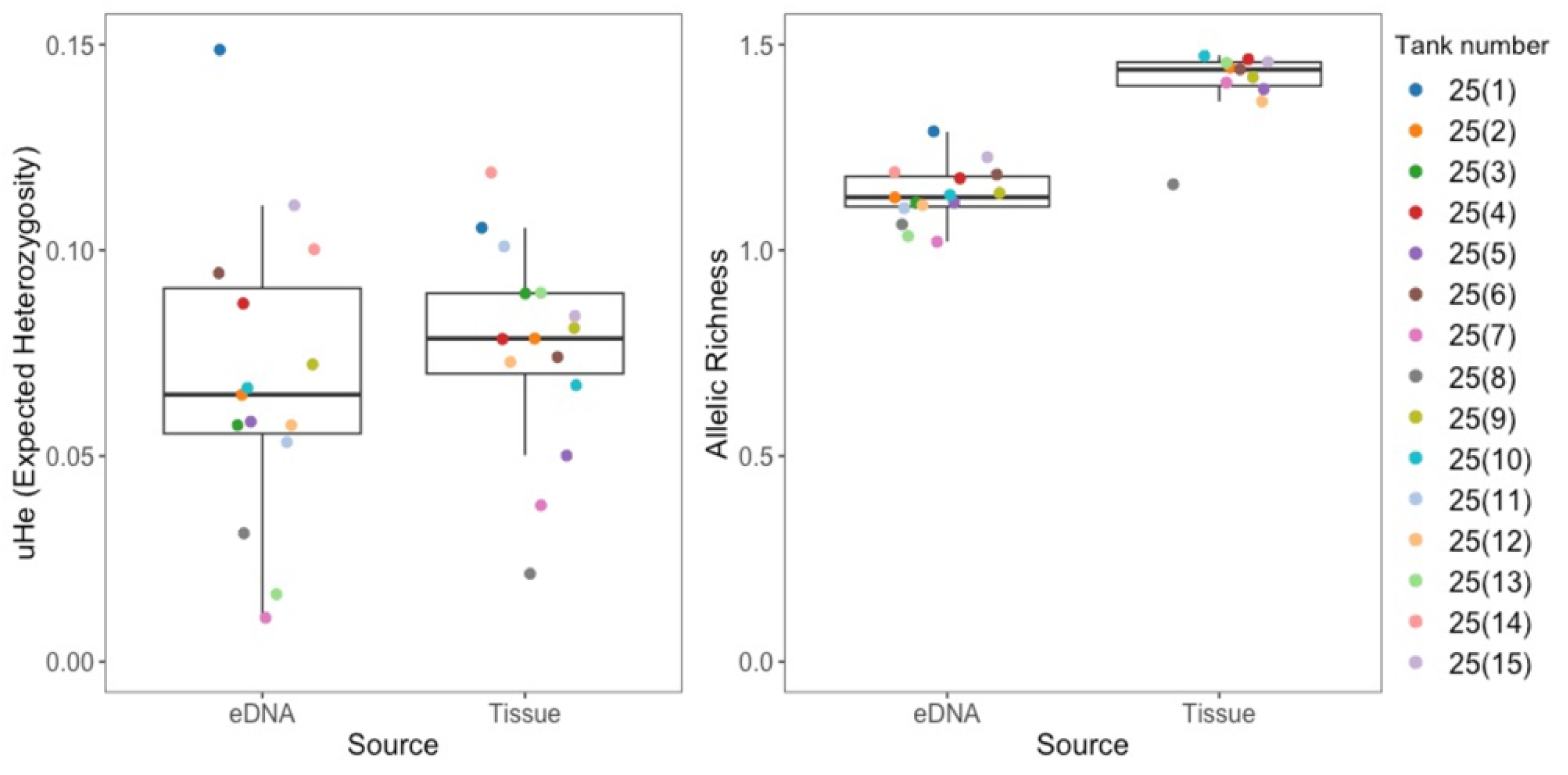
Estimates of unbiased heterozygosity (uHe) and allelic richness (AR) from eDNA samples collected from each zebrafish tank containing 25 individuals labelled as 25(1), 25(2), … 25(15) and their respective tissue samples, colour coded by tank label.

### 3.4 Contributor estimation from water samples

Contributor estimations were more accurate for eDNA samples with lower number of individual contributors of up to 20 and the negative bias between the estimated and true number of contributors increased with sample size (Figure 3). Estimation of abundance from eDNA samples containing 2, 4, 10 and 20 individual contributors had low bias of +1 to -2 individuals, with only the sample with 10 individuals showing higher negative bias of -5 individuals. Of the 15 samples containing 25 contributors that were analysed, four samples showed a bias equal to or lesser than -4 individuals, with an overall average estimation of 13.5 ± 6 individuals across all samples. In samples containing over 50 individuals, the negative bias in estimation was extremely high, with an average of -83 ± 6 individuals across samples with 100 contributors, -180 ± 11 individuals in samples with 200 contributors, -274 ± 8 individuals for samples with 300 contributors and -388 for the sample with all 411 contributors. Varying the putative number of contributors (*y*) provided to the maximum likelihood model during calculation from 50, 411 to 1000 individuals did not have any effect on the contributor estimations.

**Figure 3:**
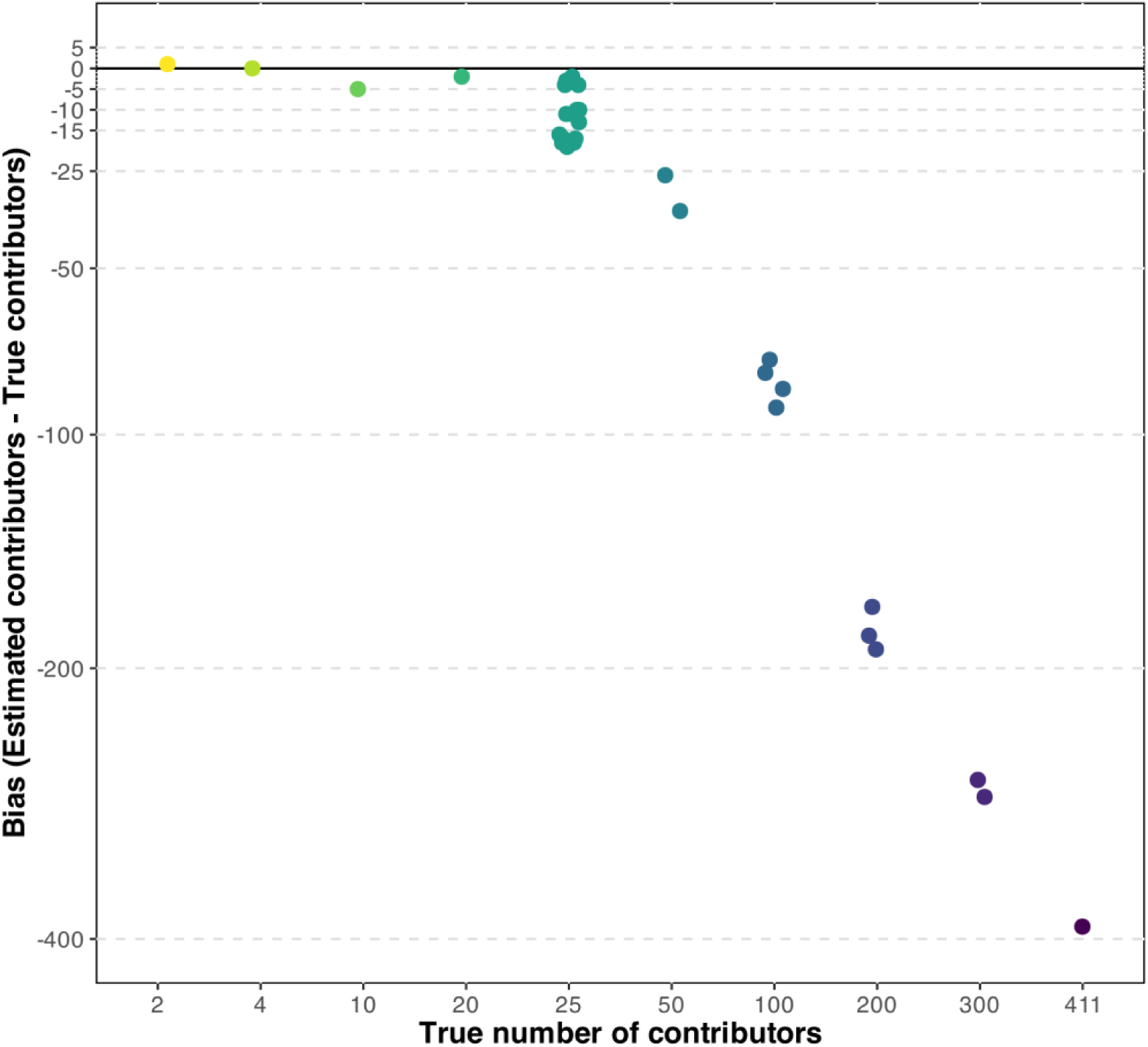
Bias in number of contributors estimated using maximum likelihood approach from eDNA samples with known number of individuals using an allele frequency threshold of 0.01. Data points are colour coded by true number of contributors.

## 4. Discussion

Knowledge on the genetic diversity, demography and abundance of a species is important in informing appropriate conservation strategies for conservation and management (Beaumont, 2007; Hohenlohe et al., 2021). Amplification of informative nuclear DNA markers from eDNA holds the potential for gaining such insight on elusive or cryptic species that could potentially live in such environments where observational monitoring is logistically challenging (Adams et al., 2019; Andres et al., 2023). This study using eDNA methods demonstrated that the amplification of a species-specific panel of nuclear microhaplotype markers can be effectively used to assess intraspecific genetic diversity, as well as estimate the number of genetic contributors in water samples containing DNA from multiple individuals. The results showed a strong correlation between population-level allele frequencies obtained from eDNA samples and those from tissue samples, validating the potential of eDNA methods for making population-level inferences about aquatic species without the need for invasive genetic sampling.

The 11 microhaplotype markers used for analysis exhibited sufficient read depths across all samples, with notably higher depths observed in eDNA samples, likely due to contributions from multiple individuals (Sigsgaard et al., 2020). The 69 SNP loci within this panel demonstrated a significantly low probability of identity, underscoring the utility of these markers for both individual identification and mixture deconvolution. The ability of microhaplotype markers to qualitatively detect individuals from complex DNA mixtures improves with higher effective number of alleles (Kidd & Speed, 2015). In the context of our study, eight out of the 11 MH markers contain five or more SNPs. This configuration likely enhances the resolution for distinguishing between multiple individual contributors within eDNA samples, thereby improving the accuracy of population-level inferences. The absence of SNPs in six out of the originally designed 17 microhaplotype markers, despite similar read depths across all 17 markers, likely results from the low genetic variation in the zebrafish sample group used for this study. All zebrafish individuals placed in experimental aquaria for purposes of this study originated from the AB strain (wild-type, ZIRC catalogue ID ZL1), bred from a single population maintained at the zebrafish facility. The lack of SNP variation could be attributed to the high degree of allele conservation within this population with limited gene flow. Therefore, the same panel of microhaplotypes may reveal more genetic variation when applied to individuals from more diverse populations.

This study demonstrated a strong correlation between allele frequencies derived from eDNA and tissue samples, both at the population level and within individual mixed samples Similar high correlations have been observed by amplifying microsatellite markers specific to round goby (*Neogobius melanostomus*) from eDNA contained in controlled mesocosm environments (Andres et al., 2021). This highlights the potential of amplifying multi-allelic nuclear markers to conduct non-invasive population genetic studies at greater resolutions than mitochondrial genomic studies, which have been the main focus of eDNA studies in the recent past (Adams et al., 2019; Foote et al., 2012). Once allele frequencies are accurately estimated from eDNA, further population genetic analyses are possible. Standard measures of intraspecific genetic variation can be obtained by evaluating allelic richness (AR) and expected heterozygosity (He), as well as assessing population structure using Principal Component Analysis (PCA). In our study, many of these analyses from eDNA exhibited high similarity to tissue-derived statistics. uHe estimates for eDNA sample with 25 individuals each were similar to those from the corresponding tissue samples but AR estimates were significantly lower, indicating a possible underrepresentation of rarer alleles and the need to optimize sequencing depth to improve detection. Additionally, estimates of uHe and AR derived from three replicates of the single eDNA sample containing all 411 contributors were lower, but without statistical significance when compared to those obtained from tissue samples and other eDNA samples. This discrepancy is likely attributed to the diluted nature of the sample, as it was created by pooling water from zebrafish tanks containing smaller numbers of individuals. From this 19 L pool, only 1 L was filtered and sequenced, likely leading to reduced read depth. The use of three replicates of eDNA samples originating from smaller groups of contributors, but collectively representing all 411 individuals, which were less diluted had population-level estimates more consistent with tissue samples.

For performing further genetic analyses such as kinship, relatedness, inbreeding, population assignment tests and so on, it is important to note that mixed eDNA samples contain a pooled mixture of DNA from various individuals and does not allow for assigning of genotypes to individuals. Furthermore, disproportionate contribution of individuals to the mixture, environmental factors of sample collection, unequal sequencing depth among samples and loci, and underrepresentation of rare alleles in eDNA samples collected in field environments can skew allele frequency estimates (Sigsgaard et al., 2020). Therefore, traditional population genetic analytical frameworks that require individual genotyping will need to be adapted towards estimating these parameters from a mixed DNA sample as well as account for the possible biases that could occur in eDNA-derived population allele frequency estimates. The theoretical and analytical frameworks existing in Pool-seq, a population genetic tool which involves sequencing a pooled mixture of DNA from multiple individuals together in a cost-effective way to infer genetic metrics without needing individual genotyping, can be adapted towards eDNA-based population studies in the future (Czech et al., 2024; Gautier et al., 2013, 2022).

Using the 69 loci analysed in this study, we were able to resolve eDNA mixtures containing up to 20 individuals with minimal bias. However, mixtures with 50 or more individuals were consistently underestimated, with the degree of negative bias increasing as the number of contributors rose. Contributor estimates were more accurate when allele frequency filtration thresholds were set at 1% or lower. In contrast, thresholds of 10% or higher led to the underestimation of the number of individuals in the mixture, likely due to the removal of rarer alleles that are more informative in estimating abundance of individuals. Furthermore, the maximum-likelihood analysis performed with greater accuracy when presented with population frequencies from the maximum possible number of individuals. Our findings were consistent with a simulation study by Andres et al. (2023), which tested the performance of the maximum-likelihood method for estimating contributors in DNA mixtures of two to 100 individuals using a panel of 64 to 256 SNP loci. The simulation, based on genetic data from a linesnout goby (*Elacatinus lori*) population (D’Aloia et al., 2020), showed that a panel of 64 SNPs could accurately resolve mixtures of up to 25 individuals. However, as the number of loci increased (from 64 to 256), the estimation of contributors in larger mixtures became distorted, potentially due to the saturation of loci across samples. The simulation demonstrated that rare alleles had the most significant impact on the accuracy of estimates, especially for larger mixtures, where the model could estimate up to 100 individuals. Thus, for improving the accuracy of contributor estimates in mixtures with a higher number of individuals using our panel of microhaplotype markers, the focus should be on enhancing the detection of rare alleles rather than increasing the number of loci. Since the zebrafish population used in this study has low genetic variation, the number of rare alleles is limited, and this method may perform better in natural populations with greater genetic variation. A similar study by Andres et al. (2021) successfully estimated the abundance of small numbers of round goby individuals (3, 5, and 10) from eDNA samples using nuclear microsatellite markers in a controlled mesocosm. These microsatellite markers also effectively resolved mixtures of up to 58 individuals in simulated DNA mixtures. Our study, using zebrafish as a model species, is the first experimental test of this maximum-likelihood model for contributor estimation on a larger scale, offering insights into its limitations.

Although this study provides robust methodology and demonstrates the use of nuclear eDNA methods to study population genetics and demography of an aquatic species, it was performed in a controlled experimental aquaria environment. Owing to this, our eDNA sequences had detection of all 69 genetic variants at sufficient read depths across all sampling replicates. However, eDNA present in natural environments pose a set of challenges such as degradation of DNA due to exposure to UV, salinity, temperature, microbial metabolism, dispersal and dilution of DNA in the water column or the presence or density of the target species in the area at the time, which could contribute to the misrepresentation of genetic variation in the population (Collins et al., 2018; Murakami et al., 2019; Tsuji et al., 2017). The low quality and concentration of the eDNA present in field samples increases the likelihood of errors that may result in the non-detection of expected alleles and rare alleles that are present at very low frequencies, ultimately leading to false negatives and an underestimation of the number of contributors to the sample and inaccurate measures of allele frequencies (Furlan et al., 2020; Sethi et al., 2019). In such cases, distinguishing between false alleles and true low-abundance alleles becomes more difficult, which further complicates the detection of rare alleles. The detection of low frequencies alleles from eDNA can be optimized by improving sample efforts, amplification, increased sequencing depth and using bioinformatics (Adams et al., 2019; Sigsgaard et al., 2020). Amplifying target markers from high-quality tissue samples (positive controls) can provide a baseline for error rates, enabling strict quality filtering (Morin et al., 2021). Retention of only the alleles found consistently in these positive controls can prevent the inclusion of errors and exclusion of true variants or rare alleles from eDNA (Andres et al., 2023).

With further development and in situ testing, the methodology outlined here could complement traditional methods for surveying the abundance and genetic diversity of invasive or endangered freshwater and marine species, supporting informed management decisions (Yoccoz, 2012). While challenges persist in estimating abundance in areas with high individual densities, this microhaplotype panel or a similar one tailored to other species, holds promise for monitoring threatened species typically found in sparse numbers (≤ 25 individuals) within sampling areas, such as cetaceans, sharks, turtles, and others (James et al., 2005; Pichler & Baker, 2000; Rosenbaum et al., 2009; Rus Hoelzel et al., 2006). This approach would be particularly valuable for genetic studies of elusive or highly migratory megafauna that span large habitats, occupy inaccessible environments, or cannot be invasively sampled due to associated risks (Dekker, 2016; Gero et al., 2014; Welsh et al., 2008). This study serves as the first step towards adapting microhaplotype nuclear DNA markers traditionally used for mixed DNA samples analysis at crime scenes towards deconvolution of mixed eDNA samples and abundance estimation of an aquatic species, with broad wide potential applications in studying the conservation genetics of natural populations.

## Acknowledgements

We acknowledge the Whadjuk Noongar people, the traditional custodians of the land on which this research work was conducted, and pay our respects to their Elders past, present and emerging. This project was supported and funded by the Minderoo Foundation through the Minderoo Foundation Exmouth Research Laboratory (MERL) and its staff. The sequencing costs of this research were partially funded the Robson & Robertson award granted by the Jock Clough Marine Foundation, UWA Oceans Institute. Housing, handling of zebrafish and sample collection for this study was conducted at the Western Australian Zebrafish Experimental Research Centre (WAZERC), Shenton Park, WA with support of the Animal Care Services staff. We would like to give our thanks to Ben Ezzy (senior technician in charge) and Wendy Hopper (Animal technician) at WAZERC for their guidance, training and support. We thank the Genomics WA Laboratory in Perth, Australia for sequencing, with special thanks to Dr. Muhammad Munir Iqbal (scientific officer) and Dr. Alka Saxena (Director) for their help and guidance. We thank Dr. Benjamin Mayne, CSIRO Oceans and Atmosphere for his advice on lab techniques and training in using laboratory equipment.

## Author contributions

Authors AM, SJ and LT designed the study. AM and LT obtained funding for the project. HP provided DArT sequence data used for MH characterization. AM undertook sample collection, laboratory work, data analysis and drafting of manuscript. All authors contributed to writing of the manuscript.

