## Supplementary material for "Towards population genetic assessments and species abundance from environmental DNA: A case study with zebrafish in controlled aquaria"

**File S1.** Link to sequence fastq files, reference MH amplicon sequences and scripts used for SNP calling and bioinformatic analysis (https://osf.io/39x6y/ )

**Table S1:** Metadata on the 17 microhaplotype markers used in this study including number of SNPs expected based on DArT sequence data versus the number of SNPs obtained from multiplex PCR of zebrafish eDNA and tissue samples used in analyses.

| Marker ID | MH sequence | MH sequence length (bp) | Forward primer (5'-3') | Reverse primer (5'-3') | No. of SNPs expected | No. of SNPs analyzed |
|---|---|---|---|---|---|---|
| **52197408** | TCCTCCATTTCGTGAGTCCAAATAAACCGAAAAAGAGCGCTTTTAACTTCTCCCCCAGCCTCCCGCTGGCCTGCAGCAGGTATACACACACACACACGCACACACAAGTGAATGCCGCTCTCTCATTGGCTGTAGGCGATGGCTGATGTTATTTTCAGTCAAAACTCATTTCACACAGTATGATTTAAATCGCCGACAGCTCTAAATATTCAGCTCGCCAAATATCTCACAGGCATTGAC | 243 | TCCTCCATTTCGTGAGTCCAA | GTCAATGCCTGTGAGATATTTGG | 3 | 8 |
| **52195134** | TGTGTTCTTCTGCTGCTGTAGCCCATCCGCCTCAAGGTTGGACGTGTTGTGTGTTCAGAGATGCTCTTCTGCAGAGATCGGTTGTAACGAGTACTTATAGTTACGGTTGCCTTTCTGTCAGCTAGAACCAGTCTGGCCATCCTCCTCTGACCTCTGGCATCAATGTGTTTAATACTGGAGCATGCCAACACGAGGAGAACA | 204 | TGTGTTCTTCTGCTGCTGTAG | TGTTCTCCTCGTGTTGGCAT | 3 | 8 |
| **52197135** | GAAACACCCACACACATACACTACAGACAATTTAGCCTACCCAATTCACCTGTACCACATGTGTTTGGACTGTGGGGGAAACCAGAGCACCCAGAGGAAACCCATGCGAAGGCAGGGAGAACATGCAAACTCCACACAGAAACGTCAACTGAGCCGAGGCTCAAACCAGCG | 173 | GAAACACCCACACACATACA | CTCAGTTGACGTTTCTGTGT | 3 | 7 |
| **52197601** | CCAAAATGCTTACACACCAGCGACTTCATTGAAAGAGGAGAGGGTTTAAGCTCTGTCGACGGGTCTCCTGCGTCTGCAGTTTAACAAGCACTTCAACAAGCCTGTTTTTTCCCGCTTGGCAAGCCAAGCTGACGTGACATGGGGGCGTGGCAGCATCGACGATTCTATTTTTTGATTCGATAATCAAAATTGAGCATAATTTTCGATCGA | 210 | CCAAAATGCTTACACACCAG | TCGATCGAAAATTATGCTCAATT | 3 | 3 |
| **52196361** | GCAGAGCCCGATAGTGGTTATCATTCATTCATTTTCTTTTCAGCTTAATCCCTTTATTAATCTGGAGTCACCACAGCGGAATGAACCGCCAACTTATCCAGCATATGTTTTACGCAGCGGATGCCCTTCCAGCCGCAACCCATCCCTGGGAAACACTACGGACAATTTAGCCAACCCAATTCACCTGTACTAC | 196 | GCAGAGCCCGATAGTGGTTA | GTAGTACAGGTGAATTGGGTTGG | 3 | 5 |
| **52197511** | TCACCAGATCCTCTCATTTAAGCCCTCGGTTCACCCGAGCTGCAGTTGTACAAATTTAGTTAATTCACACAGCAGGCTGTACTGTTTGTGTTTGGTTCACTGAATCATGCATGTGCAGTATTATCGGTTCACCGCTTCTCAATGTAACGTTGCTGTTTTAATAACAGCATAACAGTATTTATTGGATTATTTAGAGTTAGAGGAGGG | 210 | TCACCAGATCCTCTCATTTAAGC | CCCTCCTCTAACTCTAAATAATCCA | 3 | 11 |
| **52196981** | GGAGAAAGACTGACTGCATTTACATGGAAATCAGTAATCAAAATATTTGCCAAATAATTAAACAGTGAACTTTAACTACAGTTTGGCACTTTCATTCAGGAATTTCATGCATGCTCCCATGACAAACGAGATATATGATGCGAGAAACTGCTGGAAGAGTGTAGTTTTAATGGGATGTTTGATACAACACGACAAATGGGA | 204 | GGAGAAAGACTGACTGCATTTAC | TCCCATTTGTCGTGTTGTATCA | 3 | 0 |
| **52196613** | GGTAACTTGTTTGCAGTTTTGCAAAAAATTCCCTTTTCAAATTGGCTGTAAAATATAGAGGCTAAGAGAGCTTAACGTGCTGCAGTTTAAAGCCTGGTTTATACTTCTGCGTCAAGTGACCGGCATAACCCACGGCGCATGCAATACGTGTA | 152 | GGTAACTTGTTTGCAGTTTTG | CGTTTCAGAAGGGTTAATGC | 3 | 0 |

| | | | | | | |
|---|---|---|---|---|---|---|
| **52196200** | ACCCAAACCCTAAACCCAACCATACTGTAAACTTATGAATTGTTGTTTTAAAGTGTTACAAAAATGATGCTAAATTGATGGTGTATCAGCAGCTGTAATCTACCTAGACTACACAAAACAGAAATATGAATAAAACACACGAAACTATATTTGTACAAGTTAGGATTTGAACCAAAAAAGTAATAGGAAGCACAGTATCTACGTGCACACGTACAGTCCTCTAGG | 225 | ACCCAAACCCTAAACCCAACC | CCTAGAGGACTGTACGTGTGC | 3 | 5 |
| **52197653** | AGTTGTTTGAGATACGGTGACACGCAGTCAAATATTTCGCCGAACAGATCAGCCACTTTTGACGCTCAAAAACAATCAAAGCCCTTGTGCTGCAGGAATTAGGAGGCTTGCTGAAGGTGCAGCTGTCATGCAGTGAGGGGTTTGCGTCTTTAATAAACTACGTCAGTTTGCATTCACCGAACAGTAAGAA | 190 | AGTTGTTTGAGATACGGTGACA | TTCTTACTGTTCGGTGAATGC | 3 | 14 |
| **52195109** | AATCTCCAGAAATCTGCGGAATTCTGTGGAAAATCTGCAGAATTCTGCGCGCGCAGATTCCATGTGGGCCTAATAATGAAGTAAAAATAAATGCATATCATAAGATGATGAAAGTGTTTTTTTGACCTTGCATGCATTTCAGCCTGTTGTTGTTGGTA | 158 | AATCTCCAGAAATCTGCGGAAT | TACCAACAACAACAGGCTGAA | 3 | 5 |
| **52195058** | AGTGCTTGCCAAGCTTACGATTAAATTGCATATTAAGAAACAAGAAAAATTCTAAACGTTTGAGTCACTAAAAATCTGCAGAAATAGACATCAGGTCCTGGCCCCTTCCCCGAAGAATCCTCAGTCTATATAGGGATCACTGAATGGCTCTTACTAGAAGGCGG | 164 | AGTGCTTGCCAAGCTTACGA | CCGCCTTCTAGTAAGAGCCAT | 3 | 0 |
| **52197413** | CCAACAATCATGCCACATTCAAAGCTCCTTAAATCACCTTTCTTCCCCATTCTGATGCTCGATTTGAACTGCAGCAGATTGTCTTGACCATCGCTGCGTCCGAAACTGCATACTTCCATACTATATAGTCCGCTAAAATCAGTATGTGAGCCGAGTAGTATATCCGAATTCATAGAATTCAAAAATCAGTATGCGATGACTTACTACTTCCG | 212 | CCAACAATCATGCCACATTCAA | CGGAAGTAGTAAGTCATCGCATA | 4 | 0 |
| **52195232** | GCGAAACTATAAAGTGGAGTCAGCAAAAAAGCTTAACGTTTGACTGTTGCTGTTTTAAATTAGACTAAAGAAACTAAATTGTCATCCACTCTGCAGATCTCTTGCACGCCTCCGTACTGAACATAACCCCTTCCGTAATTTTTTCTTCTTGACTGATTTCTGTGAGATTCTGTAATAGTTATAGATTCGGTCATGAACATAGTAATACGTCGTGTCAATTAATATCTTGGGCAGGGAAATGTG | 243 | GCGAAACTATAAAGTGGAGTCAG | CACATTTCCCTGCCCAAGAT | 3 | 2 |
| **52196450** | CCTTTATCTACTCCTAACACGTTTCCCAACAACAGACCTCATCATTCTCAAGCTTCTGATGTCAAGACTGCAGCTCTGGACCTTTAAGTTTGACATATGAGAACTGGTTTCCAGATTGACACTTCACTTTTCAACCCATAGCCTTCCATTT | 151 | CCTTTATCTACTCCTAACACGT | AAATGGAAGGCTATGGGTTG | 4 | 1 |
| **52195252** | TGAAGTGTGGGATTGCATAATTCAAGCCCTTTTGCTGCAGATGTAATTTGCGATGTCTGTTTTTATTAATTGAATACGATGTAGTTGCTGCTAATGGCGAGTTATTCTTGAGGGCTGTGGCGGCTGTAGGCGATCAGGTTCTGCTGTCTGTTTCAAGTGCTTTTCTGAACGTCTCTGCATGCTTTTATC | 189 | TGAAGTGTGGGATTGCATAA | GATAAAAGCATGCAGAGACG | 3 | 0 |
| **52195143** | GTTCTTGCTTGTGTGTGTGTGTGTGTGTGTGTGTGTGTGTGTGTGTGTGTGTGTGTGAATGGGTGTGTGTGTGTGTGTGTGTTCAGGTGCAGTGCGTCTGCAGACACCGTTGCTCTTGCCCAGGAACAGGCGTCTGTATGAAGGCTGTGAGACGGCGTGTTTCATGGATCACAGTGGCATGCTGGTCTCTCTGCCGTATGACCTCAGGGTGAGACACACACACACACACACACACTCACTCACAGAATCAGTC | 251 | GTTCTTGCTTGTGTGTGTG | GATTCTGTGAGTGAGTGTGT | 3 | 0 |

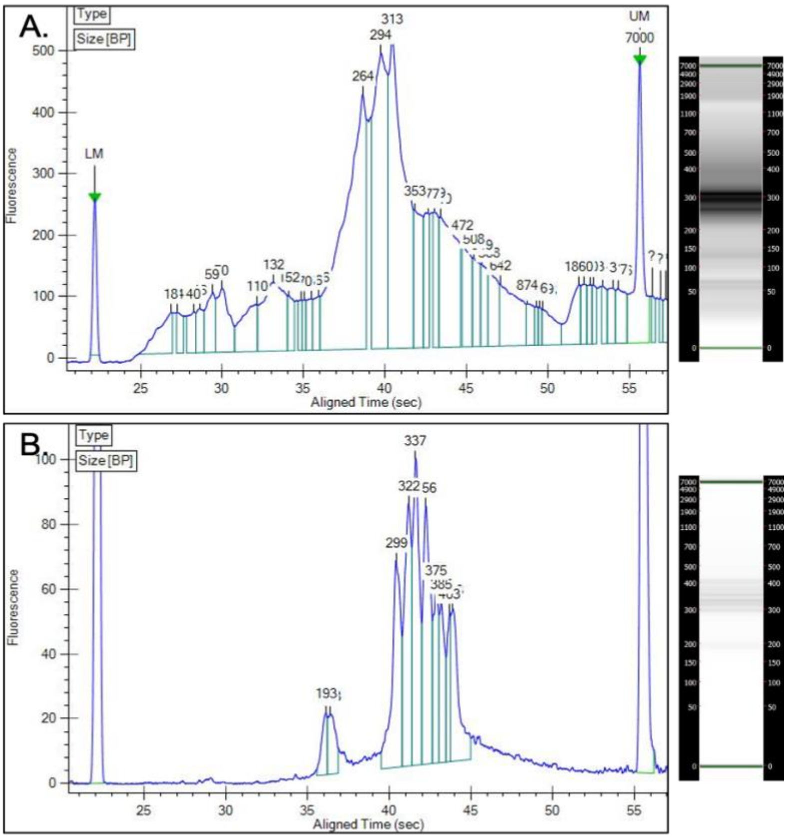


**Figure S1:** DNA fragment analysis peak table and corresponding gel image showing size (base pairs) of fragments from: A. Multiplex PCR of a zebrafish tissue DNA sample containing a panel of 40 MH markers, B. Indexed and size selected pool of 96 zebrafish samples ready to be sequenced.

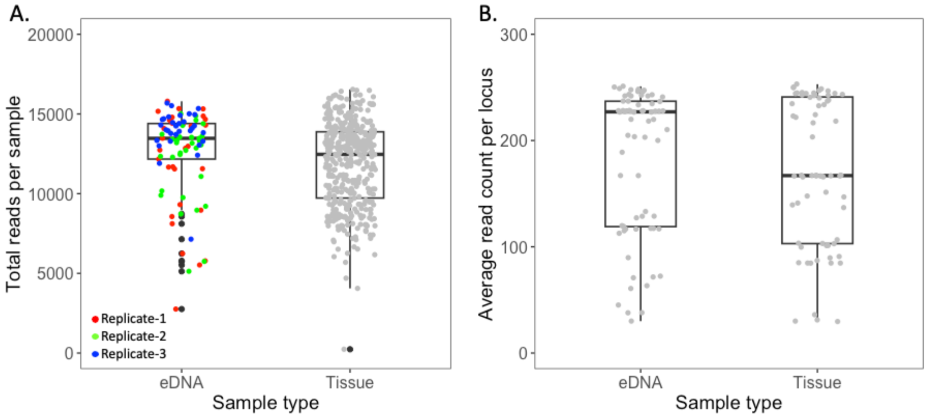


**Figure S2: (**A) Total reads per sample for each tissue sample and eDNA sample, with each eDNA replicate represented as different colours. (B) Average read count per locus across all eDNA and tissue sample.

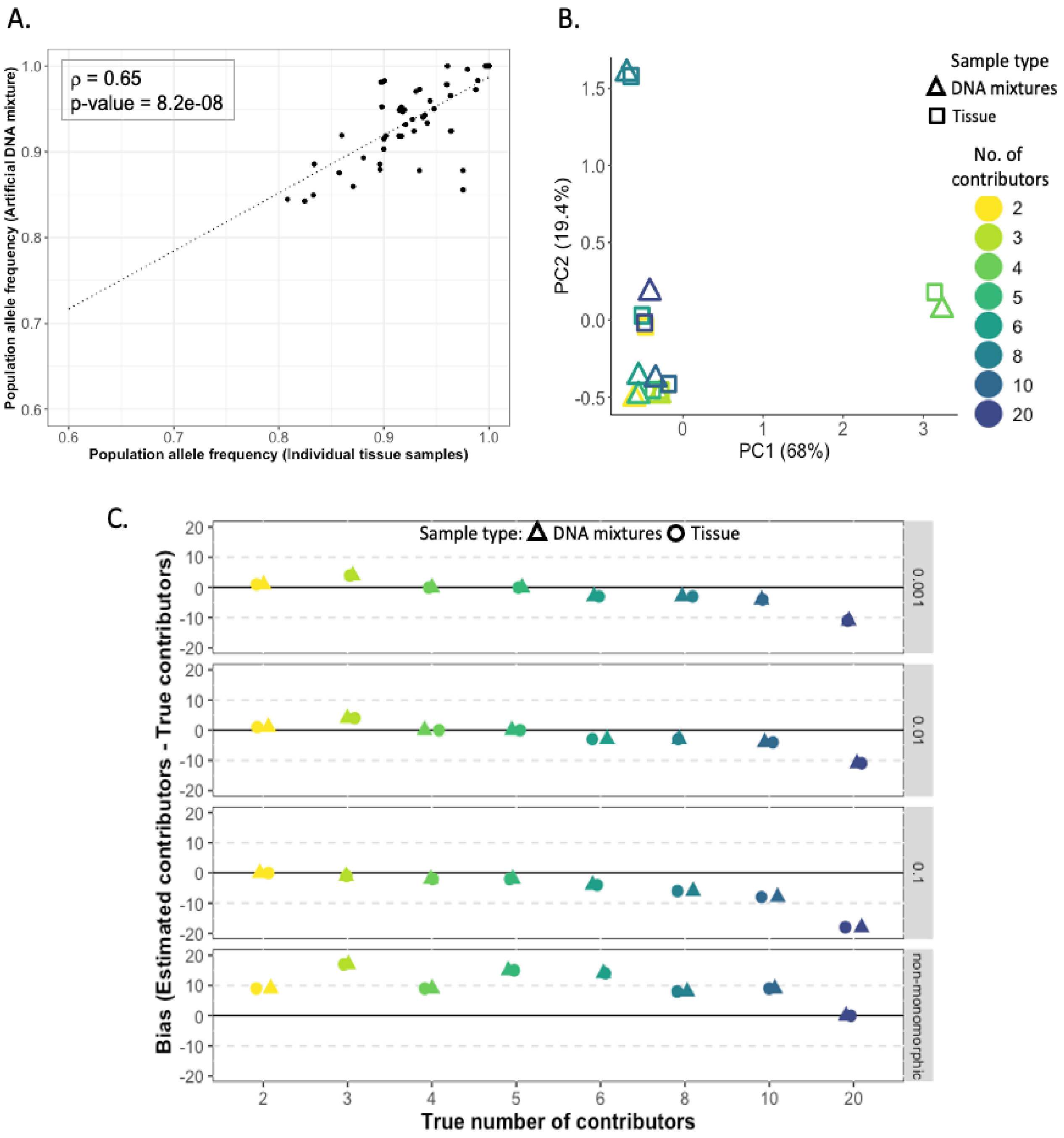


**Figure S3:** (A) Scatter plot showing the relationship between population allele frequencies for the most common allele at each SNP obtained from all artificially created pooled DNA mixtures and all corresponding individual tissue samples, with statistical significance of correlation estimated by Spearman's correlation (ρ); (B) PCA based on allele frequencies of the DNA mixtures and their corresponding individual contributors; (C) Plots showing the bias in the estimated number of contributors for DNA mixtures with known numbers of contributors calculated using population allele frequency data from 411 individuals. The triangles represent estimates from artificial pooled DNA mixtures and circles represent estimates from tissue samples. The panels indicate threshold allele frequencies below which alleles were removed (0.001, 0.01, 0.1) and a threshold where only monomorphic loci were removed (non-monomorphic).

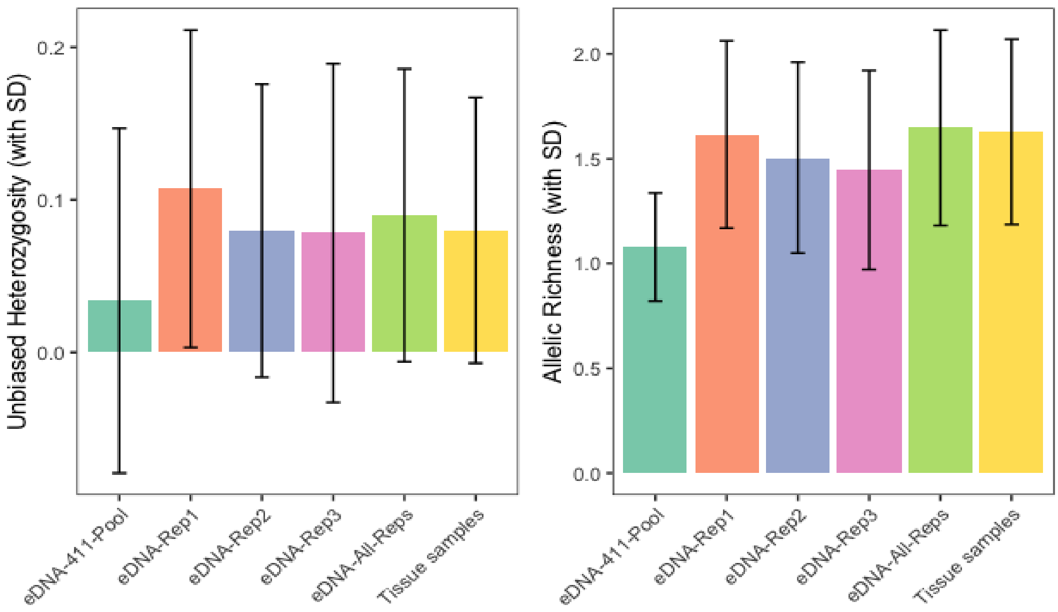


**Figure S4:** Estimates of unbiased heterozygosity (uHe) and allelic richness (AR) for the population of 411 zebrafish based on tissue and eDNA samples. Estimates were obtained from a single eDNA sample containing all 411 contributors ('eDNA_411'), as well as three replicates and pooled sequence reads from all three replicates of eDNA samples derived from smaller groups of contributors, but collectively representing all 411 individuals ('eDNA_Rep1',' eDNA_Rep2',' eDNA_Rep3',' eDNA_All Reps'). Standard deviations of estimates are represented as error bars on the graph.
